# Metabolically engineered oilcane reshapes rhizosphere microbial guilds while preserving broad functional capacity

**DOI:** 10.64898/2026.06.22.733829

**Authors:** Jaejin Lee, Baskaran Kannan, Sofia Cano-Alfanar, Hui Liu, Michael Millican, Lorien Radmer, Nicole Geerdes, Phillip de Lorimier, Bolivar Aponte Rolón, Jihoon Yang, Thanwalee Sooksa-Nguan, John Shanklin, Fredy Altpeter, Adina Howe

## Abstract

Metabolic engineering of crops can redirect host carbon flux, but its consequences for microbiomes remain unclear. Here, we show that engineering oilcane for triacylglycerol (TAG) accumulation reshapes rhizosphere microbial guilds across greenhouse and field environments while preserving functional capacity. Using 36 rhizosphere metagenomes from wild-type sugarcane and engineered oilcane accessions, we reconstructed metagenome-assembled genomes and linked community turnover with shifts in functional potential. Oilcane rhizospheres exhibited taxonomic restructuring relative to wild-type plants, driven primarily by turnover rather than nestedness and marked by genotype-dependent replacement of microbial guilds. These patterns were strongest in accession 1566 and amplified under field conditions. Despite these compositional shifts, broad patterns of functional potential remained similarly distributed, whereas pathway-level differences were evident in energy production and conversion, lipid transport and metabolism, secondary metabolite biosynthesis, transport and catabolism, and signal transduction. These findings extend evaluation of engineered crops beyond host traits alone to include microbiome-scale responses.

## Introduction

Sugarcane (*Saccharum* spp. hybrid) is cultivated on approximately 27 million hectares worldwide and contributes substantially to global sugar and biofuel production (Brant et al., 2025). Its high photosynthetic efficiency, high biomass yield, and established processing infrastructure have positioned sugarcane as a major platform for bioenergy and bioproduct development (Waclawovsky et al., 2010; Formann et al., 2020; Maitra et al., 2024). These characteristics have also motivated metabolic engineering efforts to redirect carbon storage toward triacylglycerol (TAG) accumulation in vegetative tissues. TAG has been targeted because it represents one of the most energy-dense forms of reduced carbon and can be accumulated in vegetative biomass through engineering of lipid biosynthesis and storage pathways (Cao et al., 2023).

In sugarcane, these efforts have led to the development of oilcane accessions engineered to accumulate TAG in vegetative tissues, with the goal of increasing per-hectare oil yield beyond that of traditional oilseed crops (Cao et al., 2023; Maitra et al., 2024). These accessions, including 1565, 1566, 1569, and 1580, were generated by constitutively co-expressing genes involved in TAG biosynthesis and stabilization, including diacylglycerol acyltransferase (*DGAT1-2*), oleosin (*OLE1*), and the lipid-associated transcription factor *WRINKLED1* (*WRI1*), while suppressing the TAG lipase *SUGAR-DEPENDENT1* (*SDP1*) to reduce TAG turnover (Parajuli et al., 2020).

In conjunction with plant modifications, research has also focused on the microbial communities associated with sugarcane to identify beneficial interactions. Plant-associated microbes can enhance plant nutrient acquisition, growth, and stress tolerance through mechanisms such as nitrogen fixation, phytohormone production, and improved nutrient availability. These taxa have been identified in sugarcane and other crops (Teheran-Sierra et al., 2021; Backer et al., 2018), and their manipulation represents a promising strategy to support sustainable crop production.

Previous efforts to characterize the sugarcane microbiome have shown that stable microbial taxa are present in the leaf, stem, and soil microbiomes, and that a relatively small subset of taxa can dominate total community abundance across plant compartments. In our previous study, we compared the microbiome composition of metabolically modified oilcanes with that of wild-type (WT) sugarcane (Yang et al., 2023). The 16S rRNA amplicon analysis showed differences between root-associated bacterial communities in greenhouse-grown WT and oilcane accessions, with some oilcane genotypes showing reduced representation of taxa classically associated with plant growth promotion.

In this study, we move beyond taxonomic inventories to functional characterization by analyzing 36 rhizosphere metagenomes from greenhouse- and field-grown plants. We hypothesize that oilcanes with elevated TAG accumulation host root-associated microbiomes with distinct taxonomic composition and altered functional representation relative to WT sugarcane, reflecting differences in host carbon allocation. To test this, we compared metagenome-assembled genomes (MAGs) and linked taxonomic identities to shifts in functional potential, with the goal of associating specific microbial functions with oilcane genotypes.

## Results

### Oilcane accessions show elevated TAG accumulation

Oilcane accessions showed elevated TAG accumulation under field conditions. TAG content was measured in field-grown WT and oilcane accessions 210 days after transplanting (Table 1). TAG content in leaf, stalk, and juice was significantly higher in accessions 1565, 1566, 1569, and 1580 than in WT, ranging from 103-fold higher in leaves for accession 1566 to 154-fold higher for accession 1580, and from 32-fold higher in stalks for accession 1566 to 51-fold higher for accession 1569. Root TAG content was also elevated in oilcane accessions, with accession 1580 showing a significantly higher value than WT (2.18% versus 0.13% of dry weight). Although accession 17T showed several-fold higher TAG content than WT in leaf, stalk, root, and juice samples, these differences were not statistically significant. Among all accessions, 1580 showed the highest TAG content in stem juice, reaching 1.21% of dry weight compared with 0.08% in WT (Table 1).

**Table 1.**
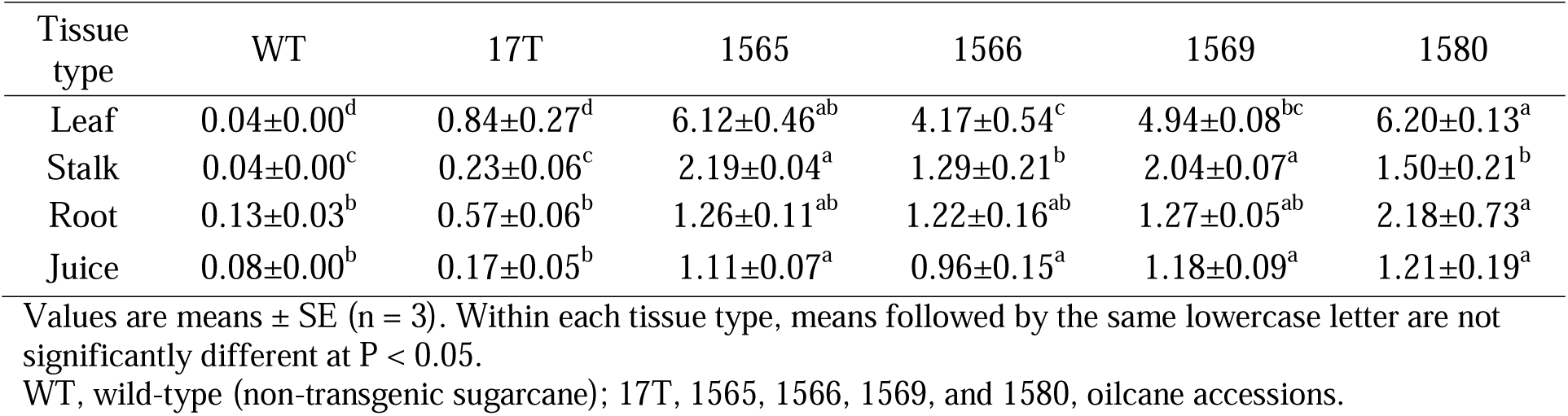
Triacylglycerol content of oilcane and wild-type plants grown under field conditions in Citra, FL.

### Transgene expression in oilcane accessions

Leaf and stalk tissues from field-grown plants were used to quantify transgene expression and target gene suppression by qRT-PCR (Table 2). Accession 17T, which carries only the *DGAT1-2* and *OLE1* expression cassettes, showed relatively high expression of these genes in both leaf and stalk tissues. In contrast, the high-TAG accessions 1565, 1566, 1569, and 1580 carried an additional *WRI1* expression cassette, in addition to the RNAi cassette targeting *SDP1*. Normalized *WRI1* expression ranged from 0.22 to 0.29 in leaf tissue and from 0.15 to 0.39 in stalk tissue across these accessions. Suppression of *SDP1* ranged from 38% to 84% in leaves and from 31% to 84% in stalks. *DGAT1-2* expression in the high-TAG accessions ranged from 0.22 to 0.77, whereas *CysOLE1* expression ranged from 0.02 to 0.11. The highest *WRI1* expression was observed in the stalk tissue of accession 1580, followed by accession 1566.

**Table 2.** qRT-PCR analysis of transgene expression and target gene suppression in oilcane and wild-type plants.

| Tissue type | Line | Expression of transgenes/suppression of target gene |  |  |  |  |
| --- | --- | --- | --- | --- | --- | --- |
|  |  | *DAGAT1-2* | *CYSOLE1* | *WRI1* | *OLE1* | *sdp1*<sup>1</sup> (% of WT) |
| Leaf | WT | 0.00±0.00<sup>c</sup> | 0.00±0.00<sup>c</sup> | 0.00±0.00<sup>b</sup> | 0.00±0.00<sup>b</sup> | 100±1.8<sup>a</sup> |
|  | 17T | 1.24±0.17<sup>a</sup> | N/A | N/A | 2.13±0.42<sup>a</sup> | 77±4.6<sup>b</sup> |
|  | 1565 | 0.29±0.03<sup>bc</sup> | 0.02±0.00<sup>b</sup> | 0.23±0.04<sup>a</sup> | N/A | 26±1.6<sup>c</sup> |
|  | 1566 | 0.48±0.09<sup>b</sup> | 0.03±0.00<sup>a</sup> | 0.22±0.03<sup>a</sup> | N/A | 16±4.4<sup>c</sup> |
|  | 1569 | 0.30±0.02<sup>bc</sup> | 0.02±0.00<sup>b</sup> | 0.29±0.03<sup>a</sup> | N/A | 62±8.6<sup>b</sup> |
|  | 1580 | 0.44±0.07<sup>b</sup> | 0.02±0.00<sup>ab</sup> | 0.27±0.04<sup>a</sup> | N/A | 18±2.9<sup>c</sup> |
| Stalk | WT | 0.00±0.00<sup>c</sup> | 0.00±0.00<sup>c</sup> | 0.00±0.00<sup>c</sup> | 0.00±0.00<sup>b</sup> | 100±9.1<sup>a</sup> |
|  | 17T | 1.38±0.47<sup>a</sup> | N/A | N/A | 2.40±0.52<sup>a</sup> | 101±11<sup>a</sup> |
|  | 1565 | 0.22±0.01<sup>bc</sup> | 0.02±0.00<sup>c</sup> | 0.15±0.01<sup>bc</sup> | N/A | 30±3.8<sup>c</sup> |
|  | 1566 | 0.54±0.10<sup>bc</sup> | 0.07±0.00<sup>b</sup> | 0.34±0.06<sup>ab</sup> | N/A | 30±11<sup>c</sup> |
|  | 1569 | 0.24±0.03<sup>bc</sup> | 0.02±0.00<sup>c</sup> | 0.21±0.02<sup>ab</sup> | N/A | 69±14<sup>b</sup> |
|  | 1580 | 0.77±0.10<sup>ab</sup> | 0.11±0.02<sup>a</sup> | 0.39±0.10<sup>a</sup> | N/A | 16±1.1<sup>c</sup> |

<sup>1</sup>Suppression of RNAi target genes are shown in % of WT. Expression of transgenes is shown relative to housekeeping gene, Glyceraldehyde-3-phosphate-dehydrogenase (GADPH).
Values are means ± SE (n = 4). Within each tissue type, means followed by the same lowercase letter within a column are not significantly different at P < 0.05.
*DGAT1-2*, diacylglycerol acyltransferase 1-2; *CysOLE1*, cysteine oleosin 1; *OLE1*, oleosin 1; *SDP1*, sugar-dependent 1; *WRI1*, wrinkled 1; WT, wild-type (non-transgenic sugarcane); 17T, 1565, 1566, 1569, and 1580, oilcane accessions; N/A, not applicable.

Agronomic measurements further showed that oilcane accessions were generally shorter, had thinner stalks, and produced lower dry biomass than WT plants under field conditions. Among the engineered accessions, 1580 showed the strongest reduction in biomass-related traits, whereas 17T produced comparatively higher biomass despite its moderate TAG accumulation. Accession 17T showed moderate TAG accumulation together with higher *DGAT1-2* and *OLE1* expression, whereas the high-oil accessions expressed *WRI1* and showed stronger *SDP1* suppression, along with substantially greater TAG accumulation and reduced plant stature and biomass (Supplementary Table 1).

### Rhizosphere MAG Composition Reveals Genotype- and Environment-Driven Community Shifts

From both greenhouse and field metagenomes, a total of 179 MAGs exceeded our minimum abundance threshold (average > 5× read coverage normalized by housekeeping gene abundances) across all samples (Figure 1, Supplementary Table 2). These MAGs were affiliated with diverse phyla, with the largest numbers belonging to Pseudomonadota (formerly Proteobacteria; n = 84) and Actinobacteria (n = 50). Within Pseudomonadota, MAGs affiliated with Rhizobiales (n = 25) and Burkholderiales (n = 13) were the most frequently assembled orders. Across all samples, the mean MAG abundance was 2.4 × 10^-4^ ± 4.2 × 10^-6^ (standard error), reported as estimated genome coverage normalized by housekeeping gene copy number.

**Figure 1.**
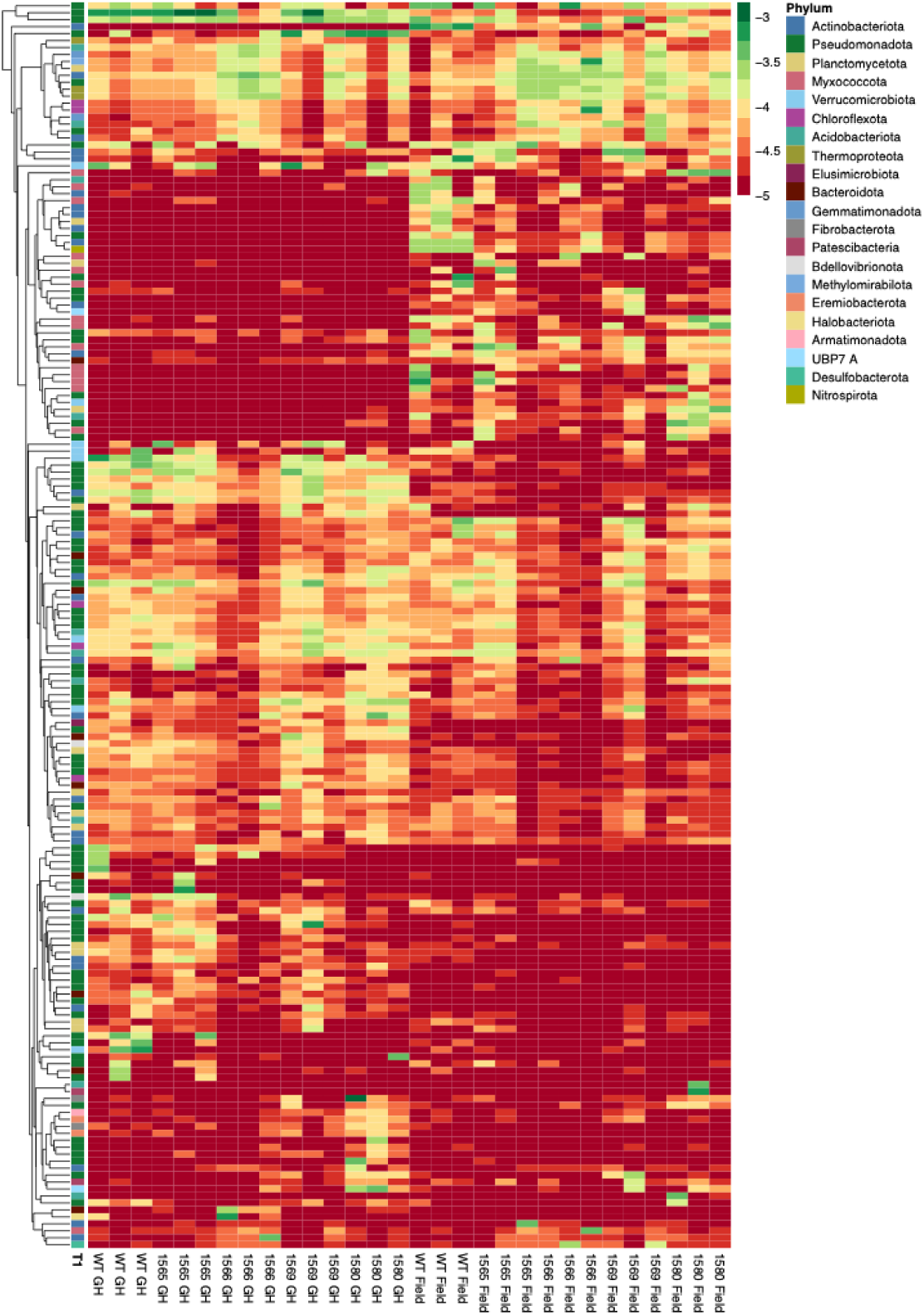
Heatmap of metagenome-assembled genome (MAG) abundances across oilcane and WT accessions and sampling times. Rows represent individual MAGs clustered by Euclidean distance, and columns correspond to triplicate samples from greenhouse (GH) and field conditions for WT (CP88-1762) and oilcane accessions (1565, 1566, 1569, 1580). Color intensity indicates normalized abundance (log scale), with warmer colors (red) representing higher abundance and cooler colors (green) representing lower abundance. MAGs are annotated by phylum (left color bar), highlighting taxonomic diversity and genotype-associated patterns in community composition.

We compared these MAG abundances in rhizosphere samples of WT and oilcane accessions across greenhouse and field environments. Ordination analyses separated greenhouse and field samples, with additional variation among accessions within each environment (Figure 2). A PERMANOVA based on Bray-Curtis dissimilarities, including accession, sampling environment, and their interaction as fixed factors, identified significant effects of accession (pseudo-F = 2.31, R^2^ = 0.223, p = 0.001) and sampling environment (pseudo-F = 6.90, R^2^ = 0.166, p = 0.001), with the full model explaining 51.8% of the total variation in community composition. Field samples generally exhibited greater variability in community composition than greenhouse samples. Across both greenhouse and field environments, oilcane samples were more like one another than WT samples, with accession 1566 consistently showing the strongest divergence from WT accessions.

**Figure 2.**
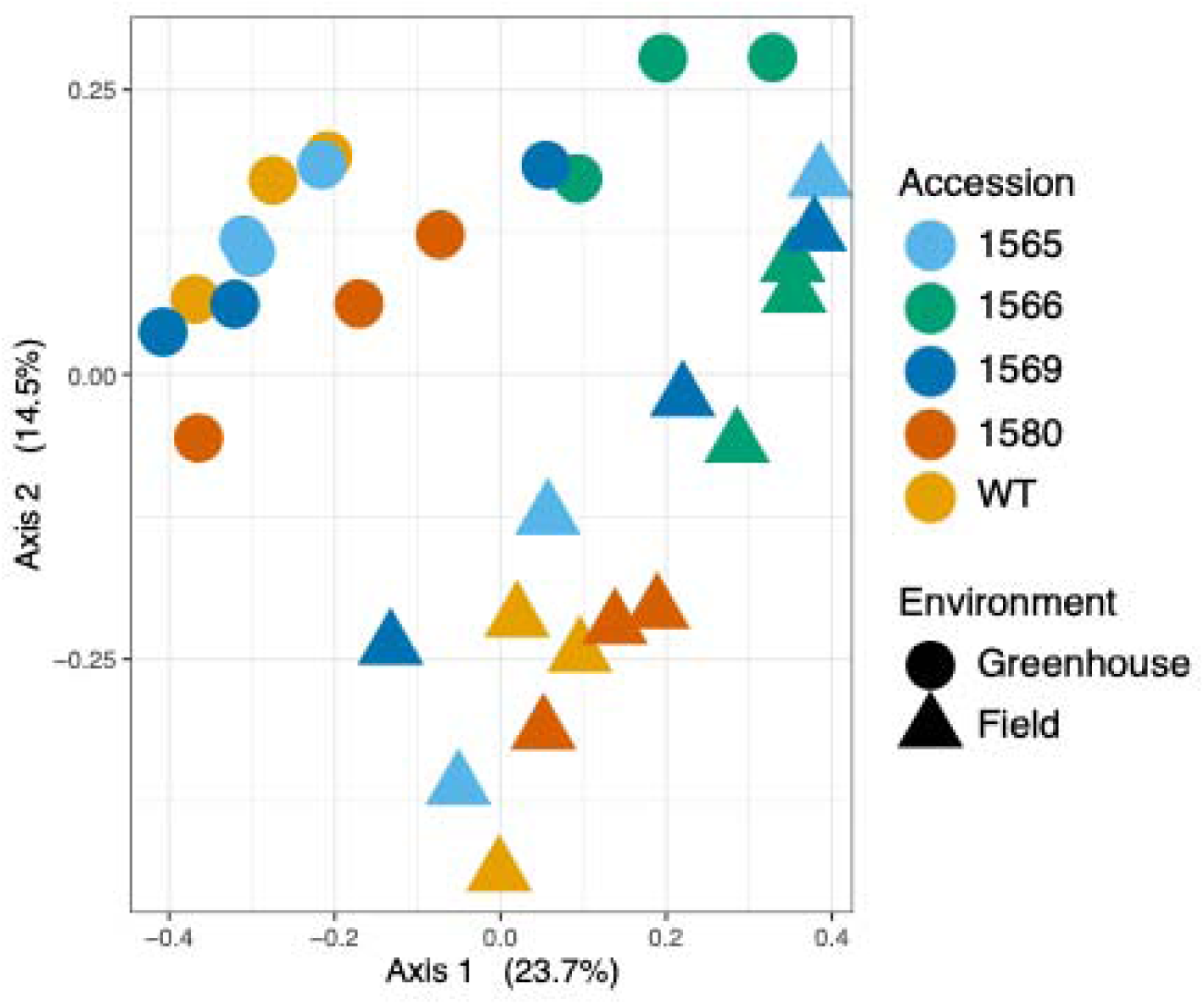
Principal Coordinates Analysis (PCoA) of rhizosphere microbial communities based on Bray–Curtis dissimilarity. Points represent individual samples colored by plant genotype (WT and oilcane accessions) and shaped by sampling time in the greenhouse (April) or field (August).

To determine whether these compositional differences reflected changes in community membership or relative abundances, we partitioned beta-diversity using the *betapart* framework, which decomposed total pairwise Sørensen dissimilarity among samples into turnover and nestedness components. Across both oilcane and WT rhizospheres, community differences were dominated by turnover rather than nestedness (Sørensen dissimilarity = 0.55 ± 0.03; Simpson dissimilarity [turnover] = 0.48 ± 0.01, nestedness-resultant dissimilarity = 0.07 ± 0.01). Similar relationships between turnover and nestedness were observed in both greenhouse and field samples.

We next identified MAGs whose abundances differed significantly between oilcane and WT accessions (Figure 3 and Supplementary Figure 3). A total of 25 MAGs met this criterion and were classified as either oilcane-enriched or WT-enriched. Oilcane-enriched MAGs were predominantly affiliated with Actinobacteria, Thermoproteota, and Chloroflexota, whereas WT-enriched MAGs were mainly associated with Pseudomonadota and Verrucomicrobiota. Oilcane-enriched MAGs were generally more abundant in field samples than in greenhouse samples, particularly in accessions 1565, 1566, and 1569 (Supplementary Figures 1 and 2). In contrast, WT-enriched MAGs did not show comparable field-level enrichment and were often similarly abundant or more abundant under greenhouse conditions.

**Figure 3.**
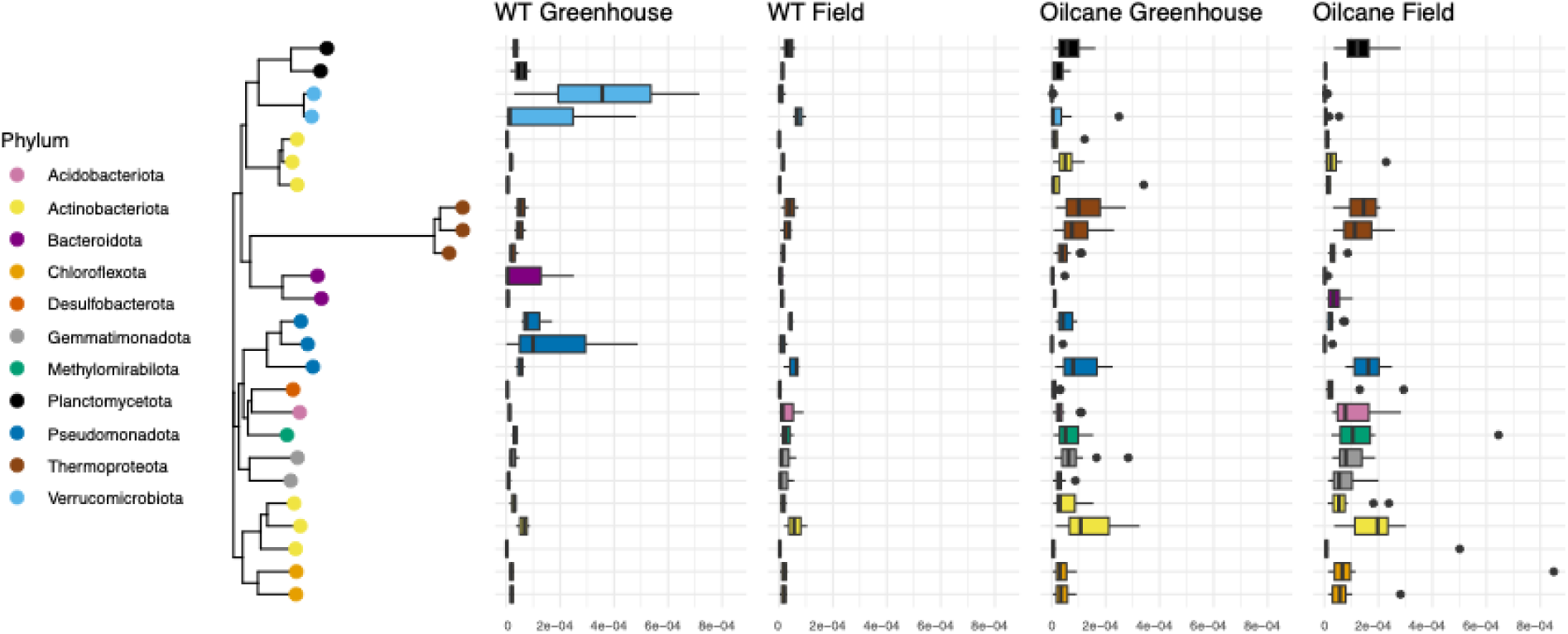
Left: Phylogenetic tree showing relationship between phyla enriched in oilcane or WT rhizosphere samples. Right: Relative abundance (normalized by housekeeping genes) of significantly enriched phyla in WT or oilcane samples taken from greenhouse (April) and field (August) conditions. Boxes represent the interquartile range (IQR) with median values; whiskers represent the range of most of the data (within 1.5 x IQR), and dots are values that fall outside beyond the 1.5 IQR.

When stratified by accession, the identities and abundances of enriched MAGs varied among oilcane lines. Accession 1565 was enriched in MAGs affiliated with Actinobacteria, Bacteroidota, and Acidobacteriota. Accession 1566 showed broader enrichment of Bacteroidota-affiliated MAGs in field samples relative to greenhouse samples. In accession 1569, MAGs affiliated with Gammatimonadota, Acidobacteriota, and Desulfobacterota were enriched under field conditions. Although MAGs enriched in either oilcane or WT were detected across multiple accessions, they were often present at low abundance. Across field samples, oilcane-enriched and WT-enriched MAGs were negatively associated (Supplementary Figure 4). This pattern was most pronounced in accession 1566 grown in the field and was not statistically significant in greenhouse samples.

### Functional Capacities Are Broadly Conserved, but Oilcane Lines Display Enhanced Metabolic and Regulatory Gene Enrichment

To assess functional potential across rhizosphere communities, open reading frames (ORFs) predicted from MAGs were mapped to Clusters of Orthologous Groups (COG) functional annotations. When expressed as relative abundances, broad functional categories showed similar distributions across oilcane and WT genotypes, although the phylum-level contributors to these categories differed between the two groups (Figure 4). When functional genes were examined as absolute abundances normalized to housekeeping gene abundances, accession 1566 exhibited generally higher gene counts than other accessions (Figure 4). Its relative phylum-level composition was similar to that of the other oilcane lines. WT rhizospheres showed broad contributions from MAGs identified as Verrucomicrobiota and Pseudomonadota across multiple functional categories, whereas oilcane rhizospheres were enriched in MAGs associated with Actinobacteria, Chloroflexota, Methylomirabilota, and Thermoproteota. In accession 1566, strong enrichment of MAGs identified as Chloroflexota and Methylomirabilota was associated with elevated absolute functional gene abundance.

**Figure 4.**
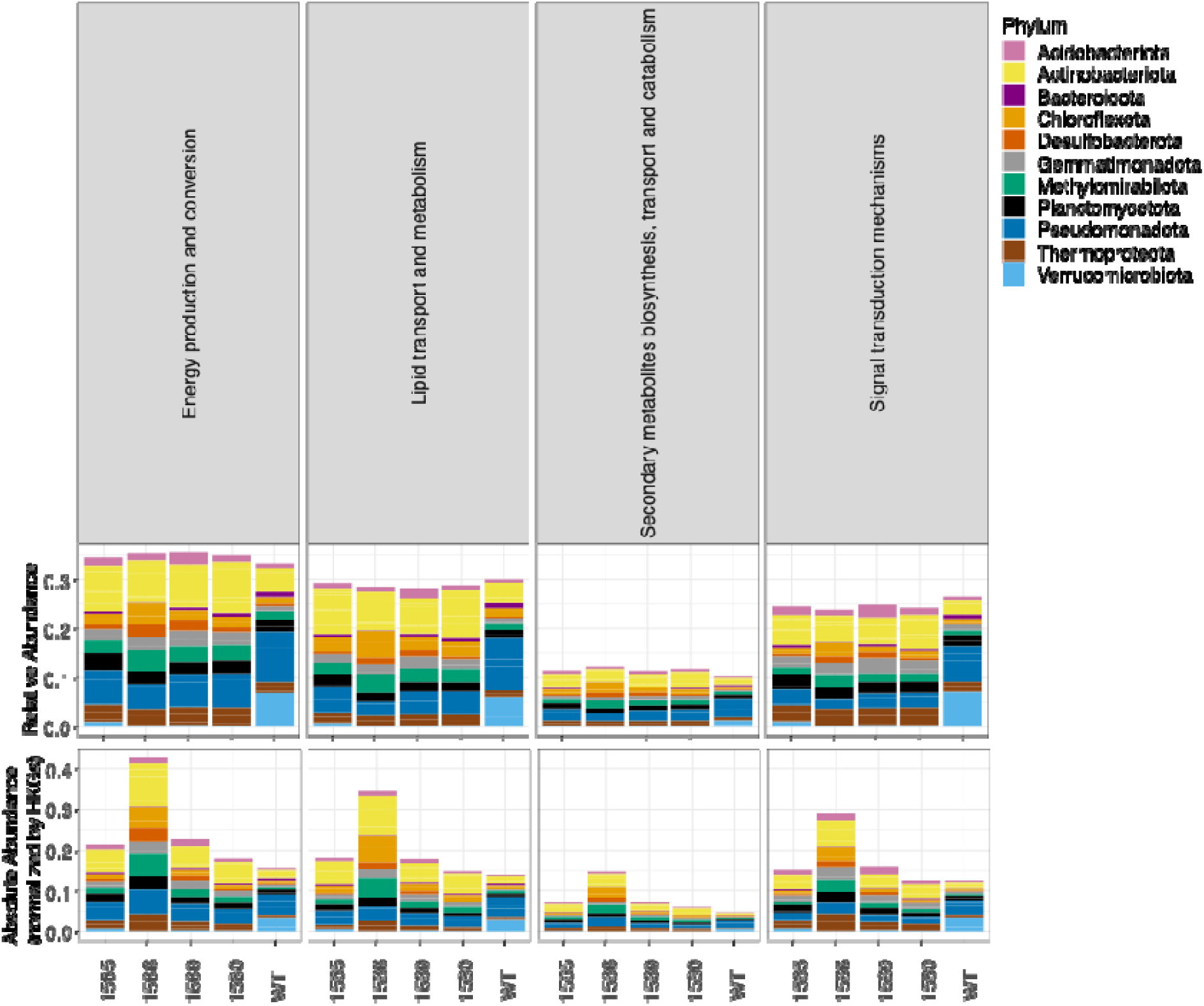
Relative and absolute (normalized by housekeeping genes, HKGs) abundances of functional classes in metagenome-assembled genomes (MAGs) in oilcane (1565, 1566, 1569, 1580) and wild-type (WT) rhizopsheres.

Across oilcane samples, genes involved in energy production and conversion, lipid transport and metabolism, secondary metabolite biosynthesis, transport and catabolism, and signal transduction mechanisms were consistently more abundant than in WT samples (Figure 4). Within energy production and conversion pathways, oilcane rhizospheres were particularly enriched in genes associated with the non-phosphorylated Entner–Doudoroff pathway, NADH dehydrogenase, the TCA cycle, and pyruvate oxidation (Supplementary Figure 5). The taxa contributing to these functions differed between oilcane and WT plant hosts, with Verrucomicrobiota contributing more prominently in WT rhizospheres, whereas Methylomirabilota, Actinobacteria, Thermoproteota, and Planctomycetota were enriched in oilcane rhizospheres. These taxa were also enriched in genes related to fatty acid biosynthesis, as well as functions associated with signal transduction and secondary metabolite biosynthesis.

## Discussion

Oilcane genotypes have successfully been engineered to redirect carbon allocation toward TAG accumulation. Field-grown oilcane accumulated significantly more TAG than WT plants, consistent with previous reports on metabolically engineered sugarcane and other biomass crops designed for lipid accumulation (Parajuli et al., 2020; Vanhercke et al., 2019a). Oilcane accessions also exhibited reduced stature, thinner stalks, and lower biomass, consistent with previously reported trade-offs associated with engineered lipid accumulation in bioenergy crops (Vanhercke et al., 2019b; Cao et al., 2023; Kannan et al., 2022). This altered host phenotype was also accompanied by reproducible shifts in rhizosphere microbial community composition across both greenhouse and field environments.

We observe that this taxonomic restructuring did not correspond to a major loss of broad functional capacity in engineered oilcanes. Beta-diversity partitioning showed that differences between oilcane and WT rhizospheres were driven primarily by turnover rather than nestedness, indicating replacement rather than simple loss or gain of taxa. Taxa enriched in oilcane and WT plants were often broadly shared across accessions, but their relative abundances covaried in a genotype-dependent manner. Broad functional categories remained comparatively stable between oilcane and WT accessions, even though the taxa contributing to these functions differed. These patterns indicate that host metabolic engineering can alter rhizosphere community assembly without collapsing community-level functional breadth and support functional redundancy among microbial contributors (Ramond et al., 2025).

When stratified by accession, the observed taxonomic differences did not reflect a uniform oilcane-versus-WT shift but instead a genotype-structured replacement of distinct MAG guilds. This restructuring was most evident under field conditions and was especially pronounced in accession 1566, which consistently diverged from WT and showed strong enrichment of Chloroflexota- and Methylomirabilota-associated functions. This pattern is consistent with our previous 16S rRNA amplicon analysis, in which accession 1566 also showed the strongest divergence from WT-associated bacterial communities (Yang et al., 2023), and with prior reports that plant genotype can shape rhizosphere bacterial community composition even under similar environmental conditions (Bulgarelli et al., 2015; Zhang et al., 2019). Most previous studies of crop microbiomes have focused on comparisons among wild relatives, domesticated accessions, or cultivar groups (Escudero-Martinez and Bulgarelli, 2023). Our results extend this framework by showing that targeted metabolic engineering of the host can also be associated with pronounced restructuring of rhizosphere microbial guilds. This suggests that microbiome assembly at the root–soil interface responds not only to natural or breeding-derived genetic differences but also to engineered shifts in host carbon allocation.

Environmental conditions represented the second strongest driver of variation in our dataset, after accession-specific effects. The greenhouse-versus-field contrast explained a large fraction of community variation, with field samples generally exhibiting greater variability and stronger enrichment patterns than greenhouse samples. This contrast is consistent with prior studies showing that sugarcane rhizosphere communities are shaped by host genotype, management regime, and environmental context (Liu et al., 2021; Moneda et al., 2022; Escudero-Martinez and Bulgarelli, 2023). Recent work has also shown that sugarcane cultivation can alter rhizosphere bacterial richness and diversity relative to wild soils, underscoring the strong environmental component of sugarcane-associated microbiome assembly (Wang et al., 2024). In our system, field conditions did not diminish host-genotype effects; instead, they appeared to amplify them, indicating that genotype-associated restructuring of the oilcane microbiome persists under more complex field settings. This persistence is notable because microbiome patterns observed under controlled conditions do not always remain detectable in the field, where environmental heterogeneity often complicates interpretation of host effects (Wagner et al., 2016; Russ et al., 2023).

Despite broad conservation of community-level functional categories, finer-scale analysis revealed differences in the absolute abundances of selected pathways between oilcane and WT rhizospheres. Oilcane-associated communities were enriched in functions related to energy production and conversion, lipid transport and metabolism, secondary metabolite biosynthesis, transport and catabolism, and signal transduction. These differences were particularly evident in accession 1566, which showed stronger representation of genes associated with the non-phosphorylated Entner-Doudoroff pathway, NADH dehydrogenase, the TCA cycle, and pyruvate oxidation. Turnover between WT- and oilcane-enriched MAGs, therefore, involved not only taxonomic replacement but also shifts in the range of metabolic pathways represented in the rhizosphere. These enrichments are consistent with reorganization of microbial metabolism under engineered host backgrounds, in which plant modification may alter nutrient availability to the soil microbiome (Ji et al., 2024). Accordingly, oilcane-enriched MAGs showed expanded representation of pathways related to central metabolism, lipid-associated functions, and regulation, alongside enrichment of metabolically versatile taxa, including MAGs affiliated with Thermoproteota and Chloroflexota (Qi et al., 2024; Reji et al., 2022; Vuillemin et al., 2024).

Oilcane- and WT-enriched MAGs differed in phylogenetic composition, highlighting the need to define the ecological roles of these lineages and the host-derived cues associated with microbiome restructuring. The observed community reorganization supports a strong association between host engineering and microbiome composition, but, by itself, does not resolve specific plant–microbe functional outcomes. Taxonomic differences occurred alongside broad conservation of community-level functional categories, suggesting that genotype-associated microbiome effects may be better understood through functional and metabolic traits rather than taxonomy alone (Louca et al., 2016; Escudero-Martinez and Bulgarelli, 2023).

Although high-oil accessions exhibited reduced stature, thinner stalks, and lower biomass than WT, our data do not distinguish whether these phenotypes arise from rhizosphere restructuring or from shifts in host physiology that secondarily alter the microbiome. Distinguishing between these possibilities will require approaches that explicitly link host carbon allocation to microbial activity. These questions can be addressed by characterizing root exudate chemistry across genotypes and linking these profiles to microbial substrate use through metabolomic and stable-isotope approaches (Zhalnina et al., 2018; Pett-Ridge and Firestone, 2017). Metatranscriptomic analyses and targeted isolate-based experiments could further test whether specific microbial guilds contribute to growth and biomass trade-offs associated with elevated TAG accumulation (Zhang et al., 2021; Vorholt et al., 2017). More broadly, these findings indicate that metabolic engineering of bioenergy crops can have substantial effects on the belowground microbiome, even when broad community-level functional capacity remains largely conserved.

## Methods

In this study, we characterized the microbiomes of four high-TAG oilcane accessions (1565, 1566, 1569, and 1580) that constitutively co-express *DGAT1-2*, *OLE1*, and the transcription factor *WRI1* while suppressing the TAG lipase *SDP1*, resulting in elevated TAG accumulation. These accessions differ from the moderately TAG-accumulating oilcane accession 17T, which co-expresses only *DGAT1-2* and *OLE1*. The microbial community structure of WT sugarcane and these oilcane accessions was described previously, and plants were grown under the same greenhouse conditions (Yang et al., 2023). After 3 months of cultivation in the greenhouse, soils loosely adhering to shaken roots were collected from each plant on April 16, 2020, and were considered rhizosphere samples.

### Transplanting of oilcane accessions to the field site

Three months after greenhouse establishment in organic soil from a south Florida sugarcane production field, oilcane accessions and WT plants were transplanted in April 2020 to the University of Florida Plant Science Research and Education Unit (PSREU) in Citra, FL (29.409006° N, −82.180473° W). The field experiment was arranged as a randomized complete block design (RCBD) with four replicates on loamy sand soil. Rows were spaced 90 cm apart and plants were spaced 60 cm apart; each accession occupied a single-row plot containing eight clonal plants per replicate.

At planting, each plant received 40 g of Osmocote® Plus fertilizer. Two weeks later, 68 kg ha^-1^ N, 23 kg ha^-1^ P, and 68 kg ha^-1^ K were applied, followed by two additional applications of the same rates at eight-week intervals. Irrigation supplied 10 mm day^-1^ for the first 2 weeks and then three times per week to ensure a minimum of 25 mm week^-1^, adjusted for rainfall during the vegetative growth phase. Weed control involved hand hoeing within rows and a mini-rototiller between rows. Insect pests (mealybugs, scales, and aphids) were managed with bifenthrin (Brigade® 2EC), imidacloprid (Admire® Pro), or sulfoxaflor (Transform™) applied at label rates, whereas orange rust was controlled with pyraclostrobin (Headline®) applied at the recommended dosage.

### Oilcane sample collection for microbiome analysis

Plant- and soil-associated samples from oilcane accessions and WT sugarcane were collected after 5 months of field growth. For each of four biological replicates, we collected leaf, stem, root, rhizosphere, and bulk soil material. One leaf sample per replicate was prepared by pooling the first dewlap leaf from three culms of a single plant, and stem samples consisted of the mature lower portion of the same three culms. Roots were detached from soil by gentle shaking onto sterile paper, cut with a sterilized pruner, and placed into individual bags. After discarding large soil aggregates, the soil loosely adhering to the roots was collected as the rhizosphere sample. A bulk soil sample was obtained by homogenizing soil from the planting site and collecting a representative subsample. All specimens were kept on ice in the field and transferred to a −80 °C freezer until further analysis.

### Real-time PCR analysis of transgene expression

To assess transgene expression and target gene suppression, we performed quantitative real-time reverse transcription PCR (qRT-PCR). Total RNA was extracted from 100 mg of first dewlap leaf tissue using TRIzol® (Invitrogen, NY, USA). One microgram of RNA was treated with RNase-Free RQ1 DNase (Promega, CA, USA) according to the manufacturer’s protocol. Five hundred nanograms of DNase-treated RNA were reverse transcribed using the High-Capacity cDNA Reverse Transcription Kit (Applied Biosystems, CA, USA). Sugarcane glyceraldehyde-3-phosphate dehydrogenase (GAPDH) primers (Iskandar et al., 2004) served as the reference gene for normalization. Primers targeting *WRI1*, *DGAT1-2*, *OLE1*, *CysOLE1*, and *SDP1* were described previously by Parajuli et al. (2020). qRT-PCR reactions were performed on a Bio-Rad CFX Connect system (Bio-Rad, Hercules, CA, USA) using SsoAdvanced™ SYBR® Green Supermix (Bio-Rad) with the following cycling conditions: 95 °C for 3 min, followed by 40 cycles of 95 °C for 10 s and 58 °C for 45 s. Relative expression levels and gene suppression ratios were calculated using the 2^-ΔΔCt^ method (Livak and Schmittgen, 2001).

### Lipid analysis of oilcane

To quantify triacylglycerol (TAG) levels, we collected 100-200 mg of the first dewlap leaf and 1-cm segments of the bottom (mature), middle, and top (immature) internodes. Roots were rinsed three times with deionized water and blotted dry on paper towels. Leaf and root tissues were freeze-dried in a lyophilizer (Labconco, MO, USA) for two days before storage at - 80°C. Stem tissues were first ground using a Retsch CryoMill (Verder Scientific, PA, USA) and then freeze-dried for three days under the same conditions. All samples were shipped on dry ice to Brookhaven National Laboratory, where lipids were extracted and TAGs were quantified by gas chromatography–mass spectrometry (GC-MS) as described previously (Parajuli et al., 2020).

### DNA extraction, library preparation, and metagenome sequencing

Rhizosphere soil samples from greenhouse- and field-grown plants were used for DNA extraction. Greenhouse rhizosphere samples were collected on April 16, 2020, and field rhizosphere samples were collected on August 4-6, 2020. DNA was extracted using the DNeasy 96 PowerSoil Pro QIAcube HT Kit and the QIAcube HT robot (Qiagen, CA, USA) according to the manufacturer’s instructions. Library preparation and sequencing were performed by the DOE Joint Genome Institute (JGI). Plate-based DNA library preparation for Illumina sequencing was performed on the PerkinElmer Scicluna NGS robotic liquid handling system using the KAPA Biosystems library preparation kit. Briefly, 1.82 ng of sample DNA was sheared to an average fragment size of 436 bp using a Covaris LE220 focused ultrasonicator. Sheared fragments were size selected by double-SPRI, end repaired, A-tailed, and ligated with Illumina-compatible sequencing adapters from IDT containing a unique molecular index barcode for each sample library. Libraries were quantified using the KAPA Biosystems next-generation sequencing library qPCR kit on a Roche LightCycler 480 instrument. Sequencing was performed on the Illumina HiSeq 2500 platform with a 2 × 151 bp indexed run configuration.

### Metagenome assembly, binning, and MAG processing

A total of 36 rhizosphere metagenomes (18 greenhouse and 18 field samples) were processed using the DOE JGI Metagenome Workflow. Co-assembly of the greenhouse and field metagenomes was performed using metaSPAdes (Nurk et al., 2017). Resulting contigs and coverage estimates were used for metagenome-assembled genome (MAG) reconstruction with MetaBAT v2.12.1 (Kang et al., 2019). MAG postprocessing included removal of scaffolds not assigned to the predominant phylum and estimation of genome completion and contamination using CheckM v1.0.12 (Parks et al., 2015). Next, GTDB-Tk v0.2.2 (Chaumeil et al., 2022) was used to assign lineage to each bin by placing them into domain-specific concatenated protein reference trees. MAGs were then dereplicated using dRep v3.20 (Olm et al., 2017), resulting in 377 MAGs.

### Functional annotation and coverage estimation

Genes in each MAG were annotated using GeneMark.hmm-2 v1.05 (Lomsadze et al., 2021). Read coverage for each gene coding region was estimated using HTSeq v2.0 (Putri et al., 2022). Housekeeping genes were identified by alignment to hidden Markov models of 71 single-copy housekeeping genes (Eren et al., 2015) using an e-value threshold of < 1e-5. Gene-level coverage was normalized by dividing the read coverage of each coding region by the average coverage of housekeeping genes within the corresponding MAG. Total MAG coverage was estimated as the average normalized coverage across gene coding regions. Coverage of functional groups, including COG categories, was calculated as the summed coverage of all genes assigned to each function.

### Statistical analyses

To evaluate the effects of accession, sampling environment, and their interaction on microbial community composition, permutational multivariate analysis of variance (PERMANOVA) was performed using the adonis2 function in the vegan R package (Oksanen et al., 2026). Analyses were based on Bray–Curtis dissimilarities calculated from normalized MAG abundance profiles, with significance assessed using 999 permutations. Differences in MAG abundance between WT and oilcane groups were assessed using the Kruskal-Wallis rank-sum test in R. MAGs were considered differentially abundant only if they met both criteria of statistical significance (P < 0.05) and mean normalized coverage > 5 in at least one comparison group.

## Accessibility to data and code

MAGs are deposited in IMG (IMG ID 3300056388). Metagenomes are available in JGI IMG (IMG IDs 1327852-1327869, 1327888-1327902, and 1327904-1327905). Code to reproduce all figures is available at https://github.com/germs-lab/PAPER_Oilcane_Rhizosphere. Datasets used in the code are available at https://figshare.com/s/7d4046715d58dab32bb3.

## Author Contributions

JL, conceptualization, data curation, formal analysis, writing – original draft;

BK & SC, propagation and molecular characterization of oilcane accessions, establishment of field trials with transgenic oilcane, collection of samples for microbiome, molecular and lipid analysis;

HL, lipid analysis with GC-MS;

MM, formal analysis, investigation, methodology, software;

LR, conceptualization, methodology, investigation;

NG, methodology, investigation, validation;

PdL, methodology, investigation, validation;

BAR, methodology, investigation, validation;

JY, conceptualization, investigation, methodology, data curation;

TS, conceptualization, investigation, methodology, data curation;

JS, coordination of lipid analysis;

FA: funding acquisition, conceptualization, writing and editing of sections of the manuscript, project administration, supervision.

AH, conceptualization, methodology, formal analysis visualization, funding acquisition, writing – original draft, project administration, supervision;

FA & BK; regulatory approval, monitoring and reporting for field trial

## Funding acknowledgements

This work was funded by the DOE Center for Advanced Bioenergy and Bioproducts Innovation (U.S. Department of Energy, Office of Science, Biological and Environmental Research Program under Award Number DE-SC0018420). Portions of the work were conducted by the U.S. Department of Energy Joint Genome Institute (https://ror.org/04xm1d337), a DOE Office of Science User Facility, is supported by the Office of Science of the U.S. Department of Energy operated under Contract No. DE-AC02-05CH11231. Any opinions, findings, and conclusions or recommendations expressed in this publication are those of the author(s) and do not necessarily reflect the views of the U.S. Department of Energy.

---

**Supplementary Table 1.** Agronomic performance of oilcane and wild-type plants grown under field conditions in Citra, FL.

| Line | Plant height (m) | Number of tillers/plant | Stalk diameter (mm) | Dry biomass yield (t/ha) | Juice volume (ml)/ 100 g stem | Total soluble solids (Brix units) |
| --- | --- | --- | --- | --- | --- | --- |
| WT | 2.82±0.02<sup>a</sup> | 9±0.20<sup>c</sup> | 20.7±0.38<sup>a</sup> | 44.6±0.72<sup>a</sup> | 50.8±2.28<sup>a</sup> | 22.2±0.11<sup>a</sup> |
| 17T | 1.90±0.02<sup>b</sup> | 14±0.33<sup>a</sup> | 16.0±0.25<sup>c</sup> | 36.1±1.89<sup>b</sup> | 44.7±2.53<sup>b</sup> | 21.7±0.20<sup>a</sup> |
| 1565 | 1.67±0.04<sup>c</sup> | 10±0.67<sup>b</sup> | 18.1±0.05<sup>b</sup> | 21.6±2.51<sup>c</sup> | 47.2±2.45<sup>ab</sup> | 21.6±0.25<sup>a</sup> |
| 1566 | 1.82±0.01<sup>bc</sup> | 8±0.17<sup>c</sup> | 15.6±0.54<sup>c</sup> | 20.8±2.38<sup>c</sup> | 48.0±0.29<sup>ab</sup> | 21.4±0.39<sup>ab</sup> |
| 1569 | 1.66±0.10<sup>c</sup> | 10±0.67<sup>b</sup> | 16.3±0.95<sup>bc</sup> | 19.2±1.44<sup>c</sup> | 51.1±0.45<sup>a</sup> | 22.1±0.37<sup>a</sup> |
| 1580 | 1.26±0.06<sup>d</sup> | 6±0.33<sup>d</sup> | 14.9±0.55<sup>c</sup> | 6.2±0.40<sup>d</sup> | 23.2±1.10<sup>c</sup> | 20.7±0.30<sup>b</sup> |

Values are means ± SE (n = 4). Within each tissue type, means followed by the same lowercase letter within a column are not significantly different at P < 0.05.
WT, wild-type (non-transgenic sugarcane); 17T, 1565, 1566, 1569, and 1580, oilcane accessions.

**Supplementary Table 2**. MAGs that exceeded minimum abundance thresholds (average greater than 5 read coverage across all samples) in greenhouse and field metagenomes. MAG approximate lineage quality (MQ = medium, HQ = high), estimation of genome completion, and contamination using CheckM v1.0.12. Lineage of MAGs was estimated using GTDB-tk v0.2.2 using the JGI/IMG workflow.

(See attached excel file)

