## Supplementary Information for "Metabolically engineered oilcane reshapes rhizosphere microbial guilds while preserving broad functional capacity"

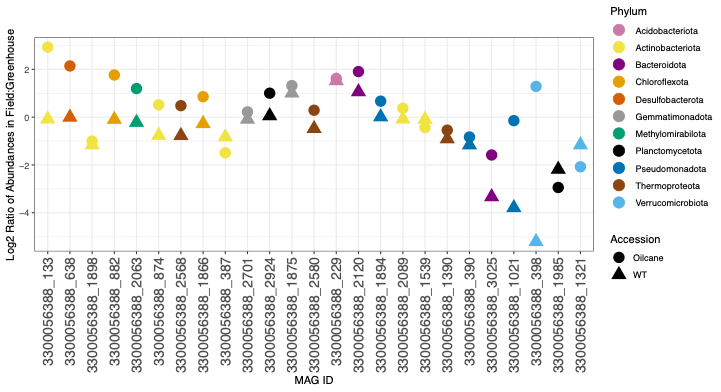

**Supp. Figure 1.** Log_2_ ratio of abundances of metagenome-assembled genomes (MAGs) between greenhouse and field conditions across WT and oilcane rhizospheres. Each point represents a MAG, colored by its assigned phylum. Positive values indicate higher abundance in field, while negative values indicate higher abundance in greenhouse.

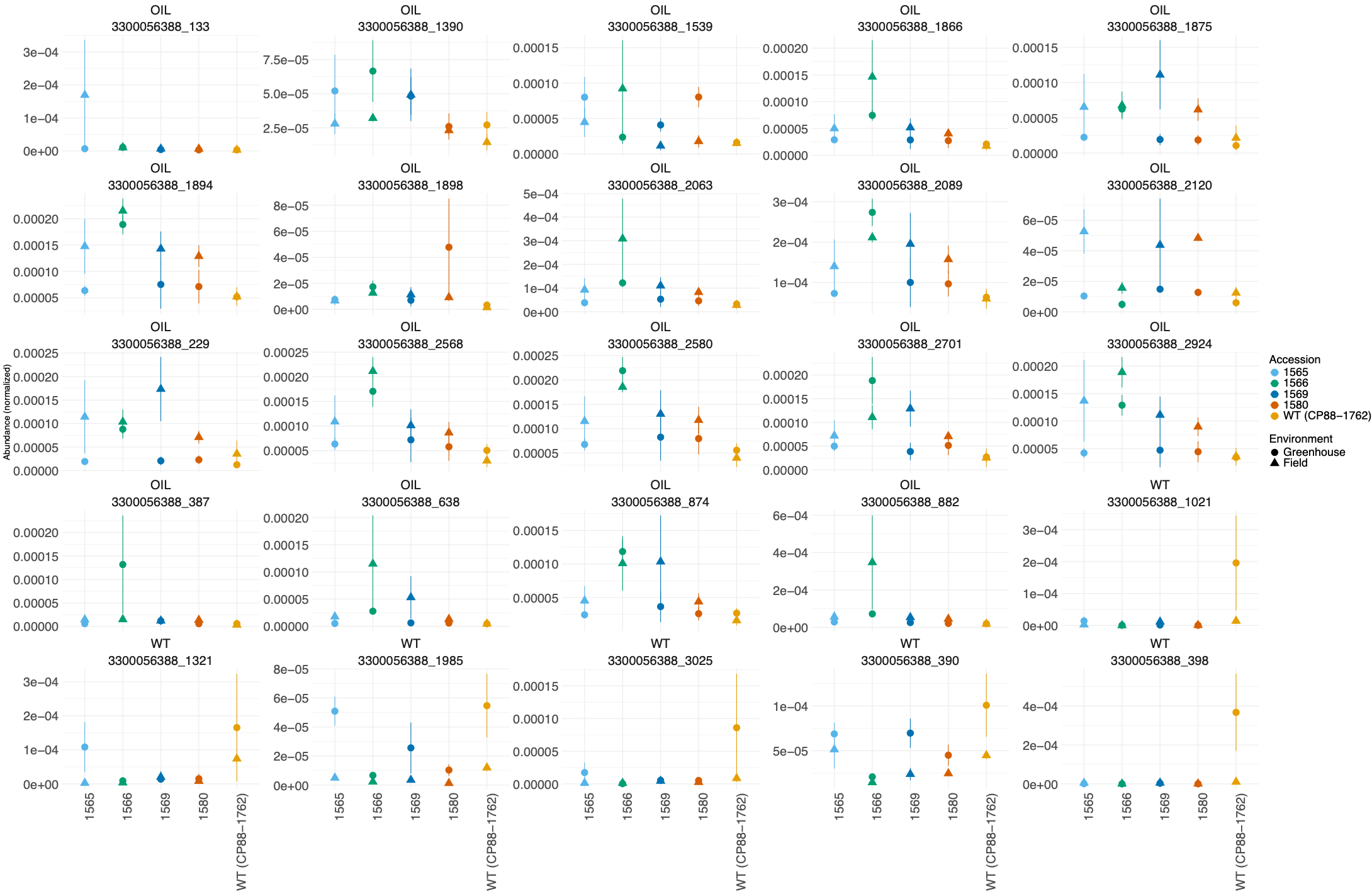

**Supp. Figure 2.** Abundances of specific MAGs in samples from oilcane (1565, 1566, 1569, 1580) and wild-type (WT) rhizosphere samples. Each MAG is classified as either enriched in oilcane (OIL) or WT. Each point represents a MAG abundance in greenhouse or field samples, and mean abundances are shown with standard error (± SEM).

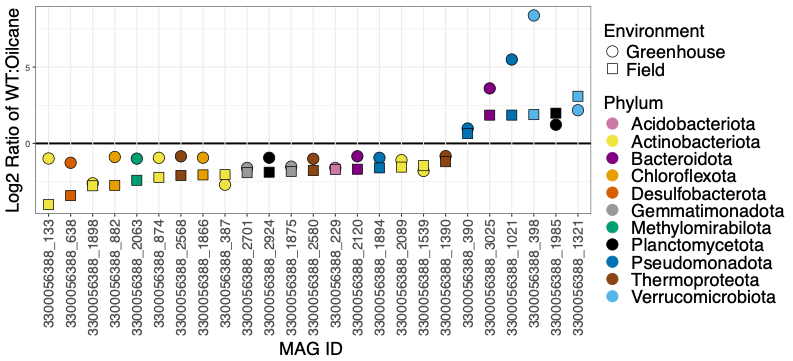

**Supp. Figure 3.** Log_2_ ratio of abundances of metagenome-assembled genomes (MAGs) between wild-type (WT) and oilcane rhizospheres across greenhouse and field conditions. Each point represents a MAG, colored by its assigned phylum. Positive values indicate higher abundance in WT, while negative values indicate higher abundance in oilcane.

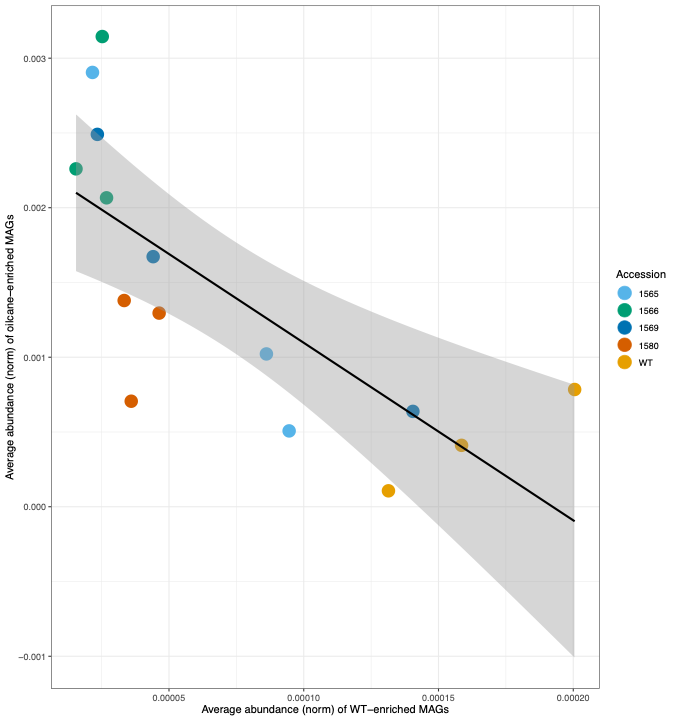

**Supp. Figure 4.** Relationship between average abundance of wild-type (WT)-enriched and oilcane-enriched MAGs under field growing conditions. Black lines indicate linear regression fits with 95% confidence intervals (gray shading). A linear regression showed a significant negative effect of the average abundance of WT-enriched MAGs on the average abundance of oilcane-enriched MAGs (β = −11.88 ± 3.01 SE, *p* = 0.0017). The model explained 54.6% of the variance (*R*² = 0.546; adjusted *R*² = 0.511). There was no significant effect found in greenhouse samples.

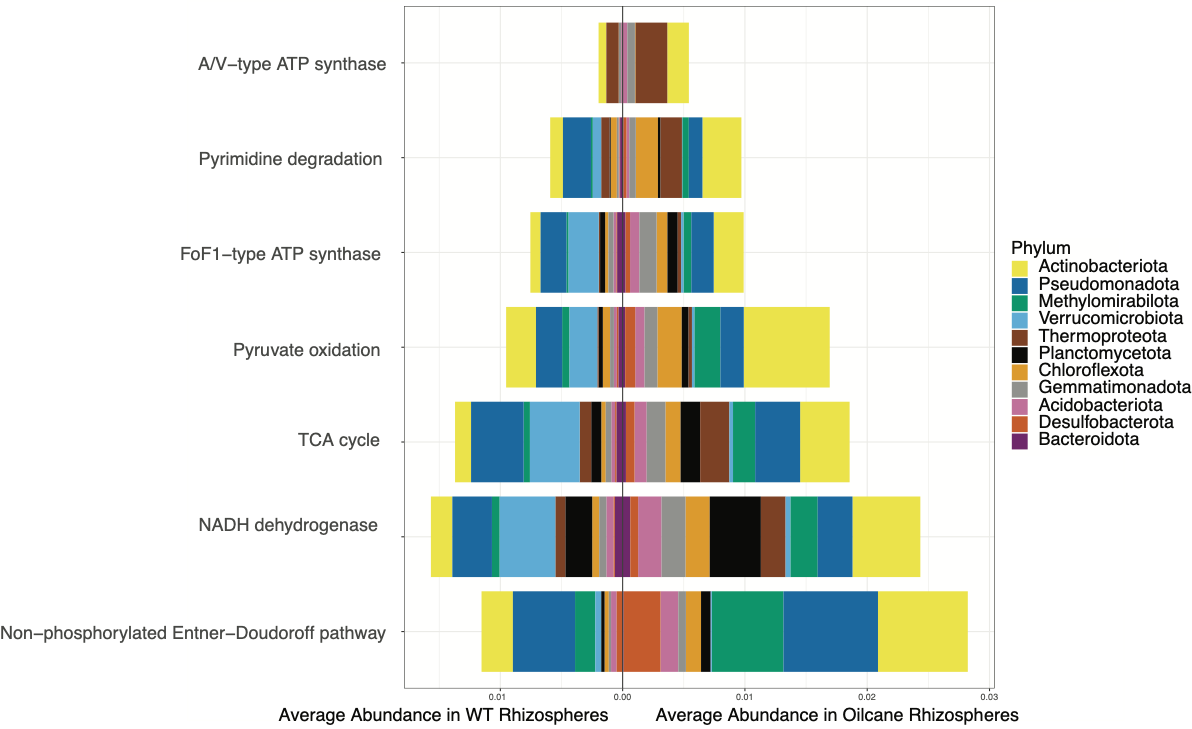

**Supp. Figure 5.** Mirror bar plot showing the average abundances (normalized by housekeeping gens) of selected metabolic pathways associated with energy production and conversion in wild-type (WT) and oilcane rhizosphere metagenome assembled genomes. Bars on the left represent wild-type (WT) rhizospheres, and bars on the right represent oilcane rhizospheres. Colors indicate taxonomic contributions at the phylum level.
